# Fluoroquinolone-Triggered Prophage Induction in *Streptococcus anginosus* Reveals Lytic Cycle, CRISPR–Prophage Interplay, and the potential for Cross-Species Horizontal Gene Transfer

**DOI:** 10.64898/2026.07.24.740447

**Authors:** Dorina Haider, Salome Barbakadze, Julia Mosler, Stefanie Mauerer, Clarissa Read, Parham Sendi, Georg Conrads, Barbara Spellerberg

## Abstract

*Streptococcus anginosus* (*S. anginosus*) has long been considered a commensal of the human microbiome but is increasingly associated with invasive infections and malignant processes. For understanding evolutionary dynamics, it is essential to investigate its mobile genetic elements, such as prophages, which are known to impact virulence, antibiotic resistance, and horizontal gene transfer. While many *S. anginosus* strains carry prophages, lysogen induction by external stimuli has not been demonstrated, and phage-mediated infection or lysis of this species has not been reported. To analyze the prevalence and diversity of prophages in *S. anginosus* genomes, we screened 140 clinical isolates by PCR revealing that 31.4% of strains were lysogenic. Correlating these findings with the presence of CRISPR immunity, we observed that *S. anginosus* strains carrying a CRISPR-Cas type II-A system were less likely to harbor prophages. Using a PCR-based approach, the spontaneous excision of several prophages of *S. anginosus* could be demonstrated and a fluoroquinolone-triggered prophage induction could successfully be established. Induction by ciprofloxacin and levofloxacin resulted in significant, concentration-dependent phage release and bacterial lysis. Transmission electron microscopy revealed viruses exhibiting the morphology characteristic of siphoviruses. Further analysis of the susceptibility of *S. anginosus* isolates and other oral and pyogenic streptococci to the isolated *S. anginosus* phages demonstrated a broad host range and the potential for cross-species horizontal gene transfer.

In conclusion, a lytic cycle of *S. anginosus* phages could be induced, highlighting their functional relevance to pathogenicity and horizontal gene transfer, while demonstrating potential clinical implications of antibiotic-mediated prophage activation.

## Introduction

Bacteriophages are the most abundant biological entities on Earth; they specifically infect bacteria and replicate within them by hijacking the host’s cellular machinery for their own reproduction (Ackermann & Prangishvili, 2012; Weinbauer, 2004). Following the selective infection of bacterial hosts, temperate phages may enter a lysogenic cycle, during which the phage genome integrates into the bacterial chromosome as a prophage and is passively replicated alongside the host genome (Edgar & Qimron, 2010; Varble et al., 2021). Prophage integration into bacterial chromosomes is widespread with 75% of bacterial isolates carrying at least one prophage (López-Leal et al., 2022). The integrated prophage is maintained in a dormant state by repressing lytic genes until environmental signals, including host stress and DNA damage, trigger the lytic cycle, leading to prophage excision (Little & Michalowski, 2010; Roberts & Roberts, 1975). While lysogenic induction can also occur spontaneously it is a rare event in the absence of external stimuli (Little & Michalowski, 2010; Lwoff, 1953). Fluoroquinolones, which are frequently used as broad-spectrum antibiotics, have been shown to induce the excision of prophages in different bacterial species (Cirz et al., 2007; Hooper, 2001; Ingrey et al., 2003; López et al., 2014). Lysogeny not only facilitates the propagation of phages but also plays a crucial role in bacterial evolution and the shaping of microbial communities (Brüssow et al., 2004; Koskella & Brockhurst, 2014). Through phage-mediated horizontal gene transfer, prophages promote the spread of accessory genes that can enhance the fitness, virulence, and environmental adaptation of bacteria (Brüssow et al., 2004; Fortier & Sekulovic, 2013; Schroven et al., 2021). Furthermore, prophages can carry and transfer genes, leading to the horizontal spread of antimicrobial resistance within microbial communities (Schroven et al., 2021; Touchon et al., 2017). Even though phages may facilitate the spread of antibiotic resistance genes, phages targeting resistant bacteria are of particular interest as promising alternative to conventional antibiotic treatment, as well as for a wide range of biotechnological applications (Bragg et al., 2014; Olawade et al., 2024; Subramanian, 2024).

Among the diverse commensal bacterial communities found in the human body, the species *Streptococcus anginosus* (*S. anginosus*) typically colonizes the oral cavity, the gastrointestinal tract, and the urogenital mucosa (Facklam, 2002; Paster et al., 2001; Pilarczyk-Zurek et al., 2022; Whiley et al., 1992). Together with the closely related species *Streptococcus constellatus* (*S. constellatus*) and *Streptococcus intermedius*, *S. anginosus* forms the *Streptococcus anginosus group* (Jensen et al., 2013; Whiley & Beighton, 1991). Although *S. anginosus* is considered a commensal, recent studies underscore its significant clinical relevance and its increasing importance as opportunistic pathogen (Pilarczyk-Zurek et al., 2022). Infections may range from mild skin infections to severe, pyogenic, and life-threatening diseases (Fazili et al., 2017). Members of the SAG are frequently associated with bacteremia, empyema and abscess formation (Clarridge et al., 2001; Laupland et al., 2018; Terzi et al., 2016; Whiley et al., 1992) and more recently, *S. anginosus* has been linked to gastric cancer (Fu et al., 2024). However, compared to other streptococci, the molecular basis of *S. anginosus* pathogenicity has scarcely been studied to date. We have previously reported on the diversity of type II CRISPR immunity and the associated metabolic burden of maintaining adaptive immunity in *S. anginosus* (Bauer et al., 2023; Haider et al., 2025). Although *S. anginosus* has been reported to carry prophages, externally triggered lysogenic induction has never been demonstrated (Brassil et al., 2020). So far, only the purified endolysin of *Streptococcus gordonii* (*S. gordonii*) phage PH15 has been reported to successfully infect *S. anginosus* (Van Der Ploeg, 2008). Therefore, this study investigates the diversity, distribution and potential impact of prophages in *S. anginosus* on bacterial pathogenicity through their excision, as well as their infectivity toward other strains of this species and related streptococci.

## Results

### Diversity and distribution of *S. anginosus* prophages

Since prophage carriage is a common phenomenon among streptococcal species, including *S. anginosus* (Brassil et al. 2020; Rezaei Javan et al. 2019), we determined the phage distribution in our clinical strain collection by PCR and PHASTEST analysis of whole-genome sequences (WGS). Due to the limited number of WGS of *S. anginosus* isolates in our collection, prophage screening with PHASTEST was restricted to a few strains. However, using this tool, prophages were predicted in the WGS of *S. anginosus* clinical isolates BSU1375, BSU1381 and BSU1701 (Wishart et al., 2023). While no prophage could be detected in *S. anginosus* SK52, the number and size of the predicted and intact prophages of the three clinical isolates are listed in Table 1.

**Table 1:** Predicted prophages in *S. anginosus* genomes and their similarity to previously characterized phages. Prophages marked with an asterisk share an identical sequence.

| Host strain | Prophage | Prophage length | Phage hit (Accession number) | Query coverage (%) | Sequence identity (%) |
| --- | --- | --- | --- | --- | --- |
| **BSU1375** | 1375_P1 | 57.8 kb | *Streptococcus anginosus* phage Javan64 (MK449000.1) | 5 | 95 |
| **BSU1375** | 1375_P2 | 46.1 kb | *Streptococcus agalactiae* phage phiStag1_20280_6_181 (PP091924.1) | 82 | 99 |
| **BSU1381** | 1381_P1* | 32.6 kb | *Streptococcus agalactiae* phage phiGBSVK-B_GBSInt9.2 (PP758847.1) | 46 | 90 |
| **BSU1701** | 1701_P1* | 32.6 kb | *Streptococcus agalactiae* phage phiGBSVK-B_GBSInt9.2 (PP758847.1) | 46 | 90 |

The NCBI nr/nt database of viruses was searched to assess the similarity of the obtained prophage sequences to previously characterized phages. Prophages 1375_P2, 1381_P1 and 1701_P1 most closely resembled annotated phages from *Streptococcus agalactiae* (*S. agalactiae*), showing high or moderate similarity, while 1375_P1 only slightly matched the previously identified *S. anginosus* phage Javan64 (Rezaei Javan et al., 2019). Notably, the prophages 1381_P1 and 1701_P1 were 100% identical. To gain a more comprehensive overview of the prevalence of prophages in a larger strain collection, the genomic DNA of 140 *S. anginosus* clinical isolates was screened for the presence of prophages previously described for this species (Brassil et al., 2020) using a PCR-based approach (Figure 1 and Figure S1). The generated PCR products in our screen were analyzed by nucleotide sequencing and following BLAST searches, excluding non-specific amplified products (Figure S1). By this approach, prophage sequences were detected in 31.4% (44/140) of the *S. anginosus* isolates. Most strains harbored one prophage (56.8%; 25/44), while some strains contained two (18.2%; 8/44) or three different prophage sequences (25%; 11/44).

**Figure 1:**
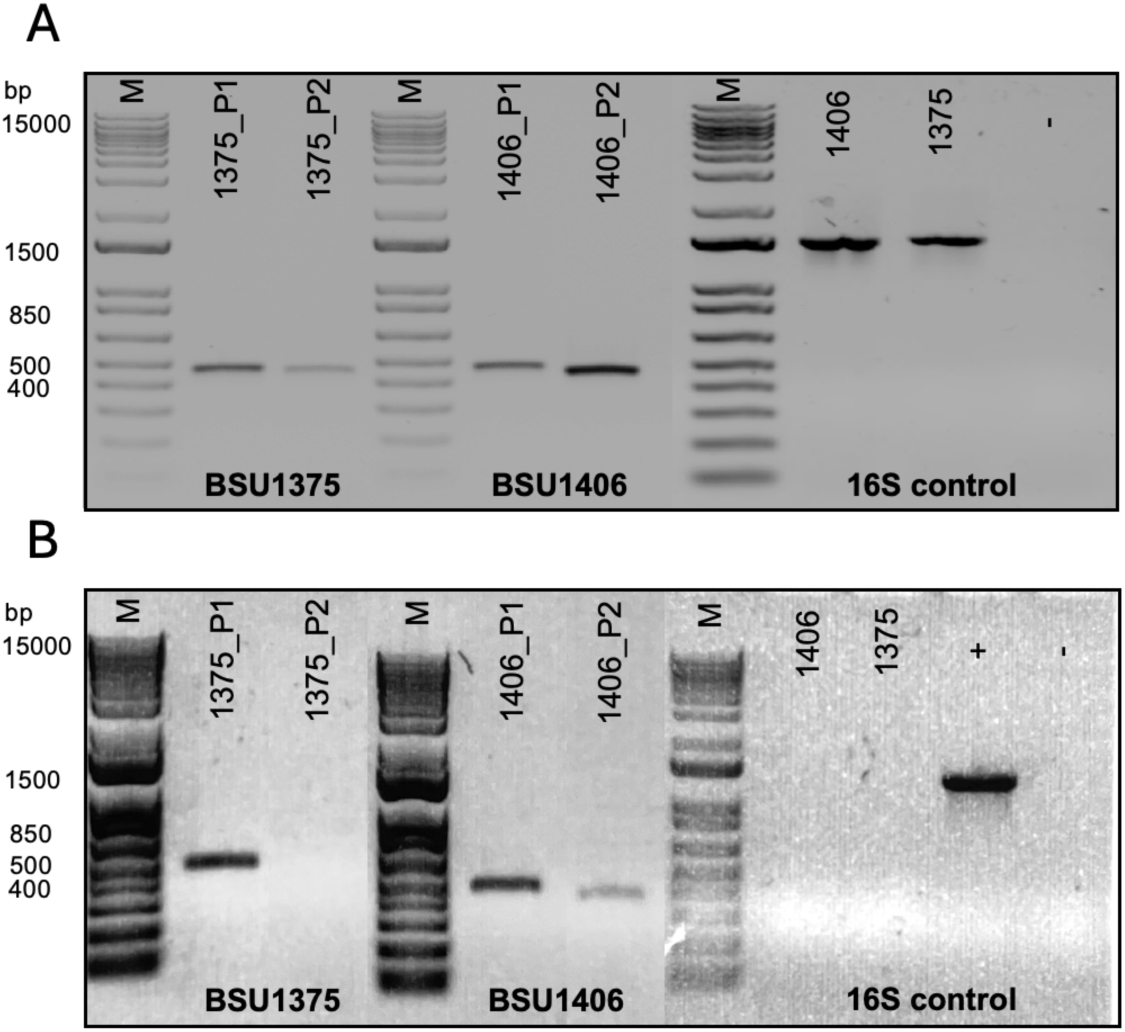
PCR-based prophage detection. A: Prophage detection in genomic DNA of *S. anginosus* isolates with published primers (Brassil et al., 2020). B: Spontaneous prophage induction in *S. anginosus* isolates. Filtered and DNase I treated supernatants of grown cultures of *S. anginosus* isolates BSU1375 and BSU1406 served as template for DNA-amplification. Specific primers were used to verify the presence of phage sequences or to identify bacterial DNA contamination (16S control). The 1 kb Plus DNA Ladder (Invitrogen) served as molecular weight marker (M). Controls are indicated by ‘+’ for positive controls with genomic DNA or by ‘-‘ for negative controls without nucleic acid.

For the species *Streptococcus pyogenes* (*S. pyogenes*), it has been shown that there is a significant difference in the number of prophages between strains with the adaptive immunity CRISPR-Cas type II-A and strains not harboring CRISPR-Cas (Yamada et al. 2019). The presence of CRISPR-Cas type II-A systems was investigated in relation to the prophage content in *S. anginosus*. Among the 140 strains of our collection, statistical analysis by a Chi-square test of independence found a value of *p*<0.05 demonstrating that in strains carrying CRISPR-Cas type II-A, the presence of one or more prophages was significantly less likely (Table 2).

**Table 2:** Comparison of prophage presence in CRISPR-Cas type II-A carrying and non-carrying *S. anginosus* strains.

|  | CRISPR-Cas type II-A presence<br>(n=82) | CRISPR-Cas type II-A absence<br>(n=58) |
| --- | --- | --- |
| Prophage presence | 22% | 44.8% |
| Prophage absence | 78% | 55.2% |

### Spontaneous and fluoroquinolone-triggered induction of *S. anginosus* prophage

Spontaneous induction of prophages integrated into the bacterial genome has been described for many streptococcal species (Baldwin & McKay, 1987; Brassil et al., 2020; Carrolo et al., 2010). The spontaneous induction of prophages could be observed under laboratory conditions in liquid cultures of *S. anginosus* strains BSU1375 and BSU1406 (Figure 1).

Viral sequences were identified by PCR analysis of filtered and DNase I-treated supernatants of grown streptococcal cultures. Nucleotide sequences of prophages 1406_P1, 1406_P2 and 1375_P1 could be detected, confirming spontaneous induction, but not for prophage 1375_P2.

For the induction of prophages with external stimuli, fluoroquinolones are used at subinhibitory concentrations; thus, the minimum inhibitory concentration (MIC) for these substances was determined in *S. anginosus* prior to phage induction experiments. The MIC was 1 µg/ml for all strains for CPX and for LVX as 1 µg/ml for *S. anginosus* SK52 and 0.75 µg/ml for the three clinical isolates. Therefore, a uniform subinhibitory concentration of 0.625 µg/ml was chosen for both antibiotics in further experiments. PCR-based analysis showed that after treatment with CPX or LVX, the number of released phages for *S. anginosus* BSU1375 increased compared to spontaneous induction, with the effect being most pronounced at the two lower template quantities (Figure 2). These data confirm lysogenic prophage induction in *S. anginosus* by both fluoroquinolones.

**Figure 2:**
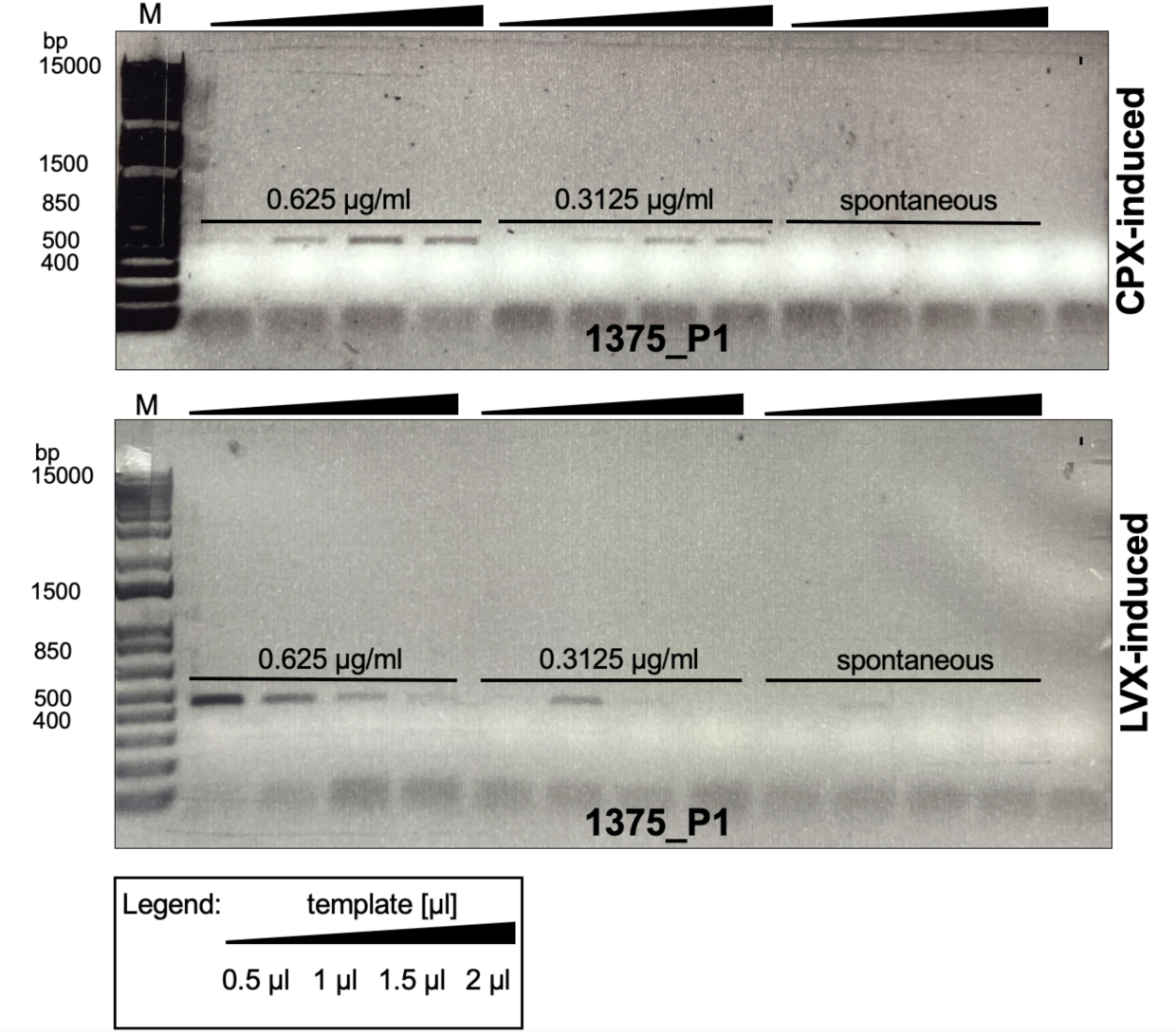
Fluoroquinolone-triggered prophage induction in *S. anginosus*. Using a PCR-based approach, filtered supernatants of *S. anginosus* clinical isolate BSU1375 were examined for phage sequences after treatment with subinhibitory concentrations of fluoroquinolones (0.3125 µg/ml or 0.625 µg/ml) or without treatment as a control for spontaneous induction. Results are shown for ciprofloxacin (CPX) and levofloxacin (LVX). Gradient shown above indicates the amount of template used (0.5 µl, 1 µl, 1.5 µl and 2 µl). The 1 kb Plus DNA Ladder (Invitrogen) served as molecular weight marker (M). Controls are indicated by ‘-‘ for negative controls without nucleic acid.

Further, the growth kinetics of *S. anginosus* isolates BSU1375, BSU1381 and BSU1701 were monitored in the presence of prophage-inducing fluoroquinolones and compared with *S. anginosus* strain SK52, which does not carry prophages. In contrast to SK52, growth of all isolates harboring prophages was inhibited after treatment with CPX (Figure 3) and LVX (Figure S2) in a concentration-dependent manner. The observed growth patterns indicate that prophage induction and subsequent bacterial lysis are effects of the fluoroquinolone treatment.

**Figure 3:**
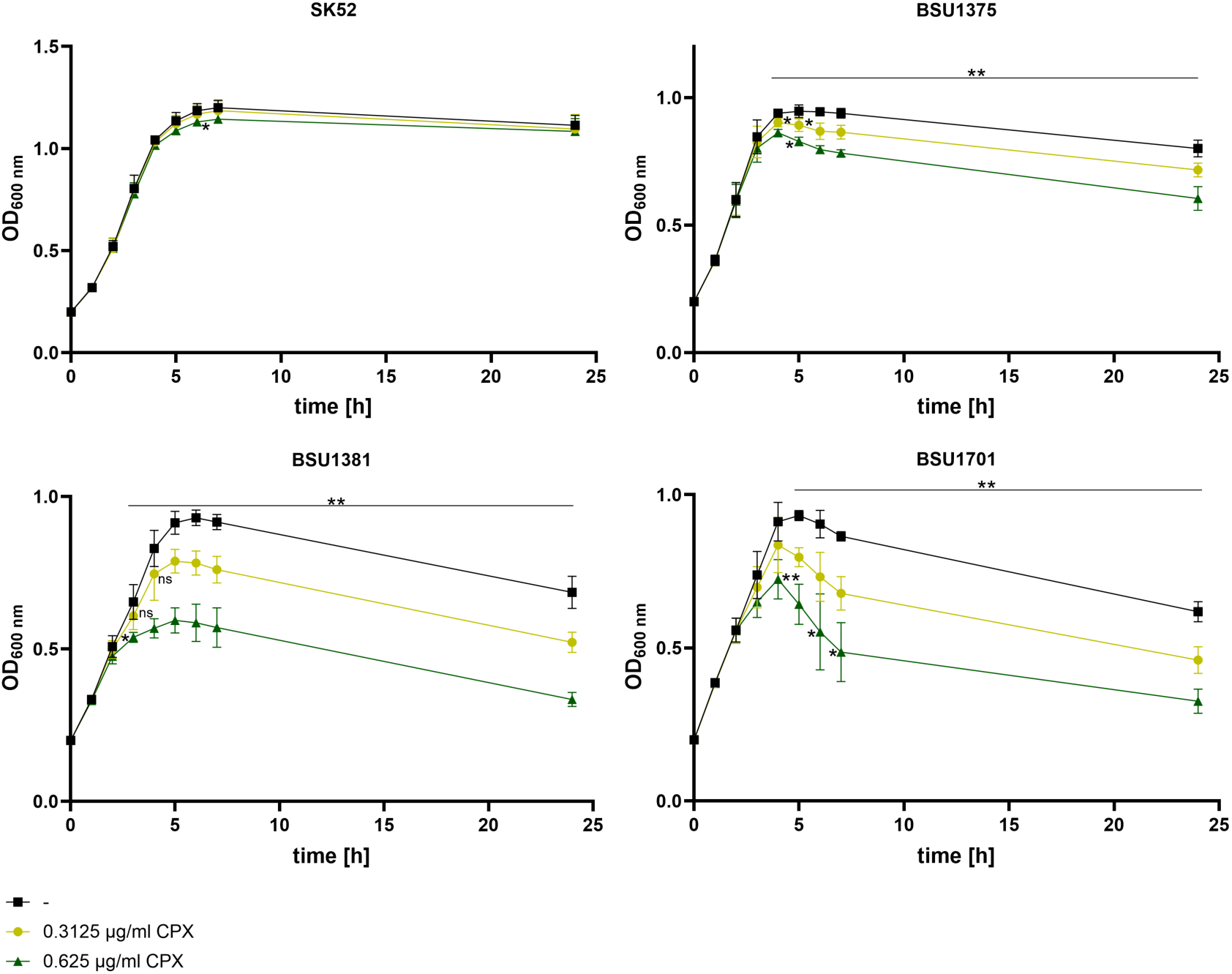
Fluoroquinolone-triggered lysogenic growth patterns of *S. anginosus* isolates. Growth kinetics of *S. anginosus* SK52 and clinical isolates BSU1375, BSU1381 and BSU1701 measured after treatment with CPX. Black curves indicate growth without treatment, while 0.625 µg/ml and 0.3125 µg/ml CPX is colored in dark and light green, respectively. Each data point represents an average of five independent measurements including standard deviation. Differences were statistically significant (*p<0.05, **p<0.01, ***p<0.001; Mann-Whitney-U test) and are shown above unless otherwise indicated. When indicated on the left side of the data point it refers to 0.3125 µg/ml and on right side it refers to the untreated control.

### Visualization of *S. anginosus* phages

Siphoviruses are tailed phages with double-stranded DNA (*Caudoviricetes*), that are abundant in streptococcal species and have also been reported in *S. anginosus* (Brassil et al., 2020; Kovacec et al., 2024; McShan et al., 2019). To visualize the morphology and structural features of the phages detected by PCR, transmission electron microscopy (TEM) was conducted. Fluoroquinolone-induced phages from *S. anginosus* isolates BSU1375, BSU1381 and BSU1701 were concentrated and visualized in multiple samples and varying degrees of intactness. The phages isolated from *S. anginosus* BSU1375 and BSU1381 (Figure 4A-E) displayed long, non-contractile tails, consistent with the morphology of *Caudoviricetes*. However, the phages from isolate BSU1701 could not be kept intact during the isolation and concentration process (Figure 4F). All phages shared the characteristic isometric head with a diameter of approximately 56 nm and a flexible, non-contractile tail of approximately 209 nm (Table 3), as described for siphoviruses (Ackermann, 2007). Tail dimensions of phages induced from isolates BSU1381 and BSU1701 were not clearly determinable due to overlaps with other structures or damaged tails, therefore only completely visible and intact capsids were considered for calculations. For some viral particles the base plate could be visualized (Figure 4).

**Figure 4:**
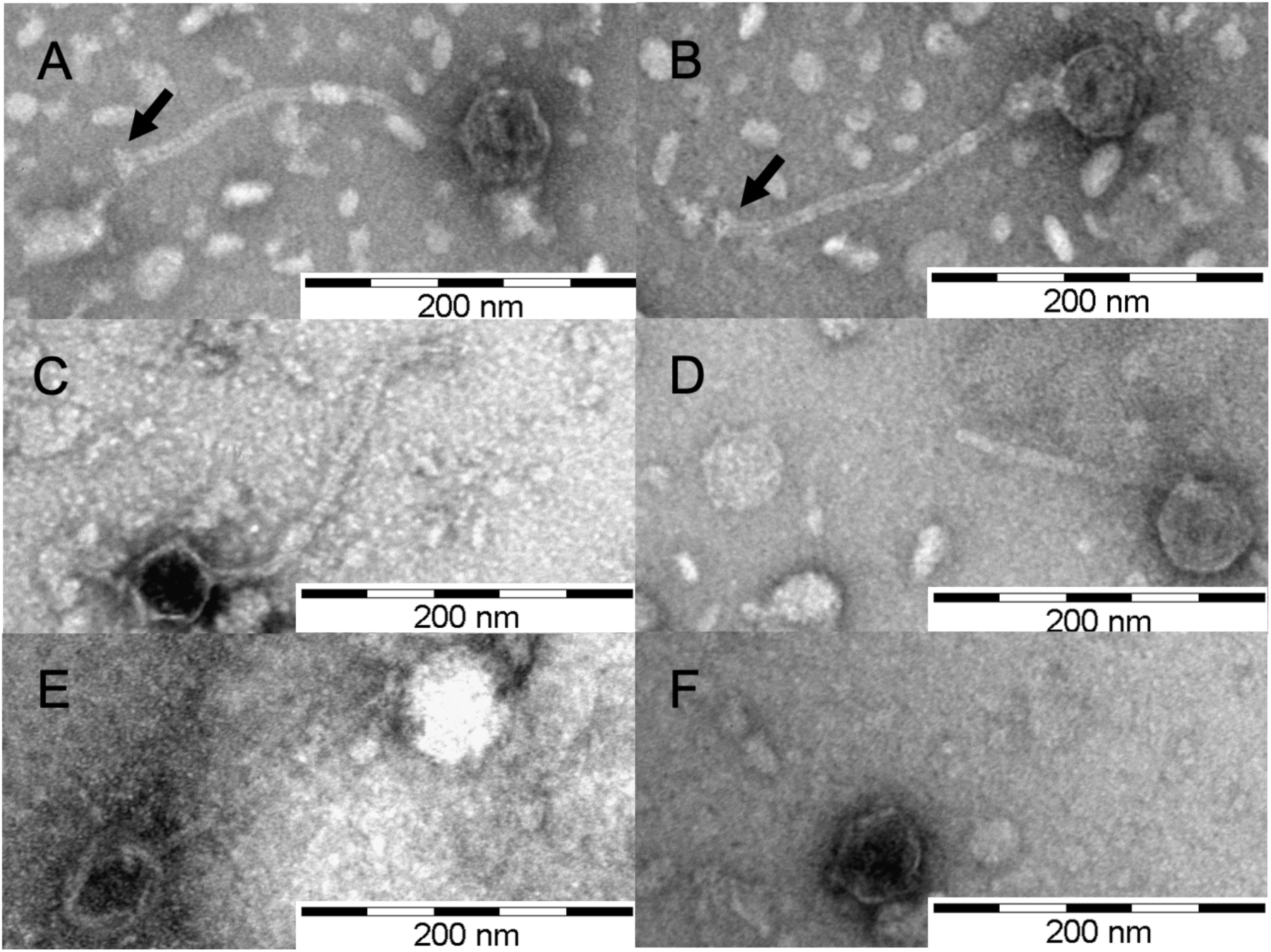
Lysogenic phages induced with fluoroquinolones from *S. anginosus* visualized with transmission electron microscopy. (A) – (C) isolated from BSU1375, partially with visible base plate (black arrow); (D) and (E) induced from BSU1381; (F) isolated from BSU1701. Scale bars represent 200 nm.

**Table 3:** Dimensions of phages induced from *S. anginosus*. Average dimensions based on a minimum of four phages.

| Phages induced from *S. anginosus* isolate | # of phages | Head diameter [nm] | Tail length [nm] |
| --- | --- | --- | --- |
| BSU1375 | 21 | $56.2 \pm 2.1$ | $208.5 \pm 14.2$ |
| BSU1381 | 4 | $56 \pm 3.5$ | not measurable |
| BSU1701 | 9 | $55.9 \pm 1.5$ | not measurable |

### Host interaction of *S. anginosus* phages

Phages possess a narrow host range, often limited to specific strains or closely related species (Brock 1964; Szymczak et al. 2019). For *S. anginosus* whole phage particles capable of infecting this species and propagating through the lytic cycle have not yet been identified. The infectivity of the phages induced by fluoroquinolones was investigated by testing the sensitivity of various *S. anginosus* strains (Table 4). Of the *S. anginosus* strains tested, five isolates were sensitive to the phages 1375_P1 and 1375_P2 from *S. anginosus* BSU1375, while one isolate each was susceptible to the phages 1381_P1 and 1701_P1 from strains BSU1381 and BSU1701. To assess whether a wider host range of streptococcal species is susceptible to the isolated *S. anginosus* phages, selected oral and pyogenic streptococci were included in the analysis. Although most oral and pyogenic streptococci tested were resistant to the applied phages, one *Streptococcus oralis*, one *S. constellatus,* and two *S. agalactiae* isolates were found to be sensitive to at least one of the phages. Phage titers could not be determined, but a concentration-dependent clearance of the bacterial lawn was observed.

**Table 4:**
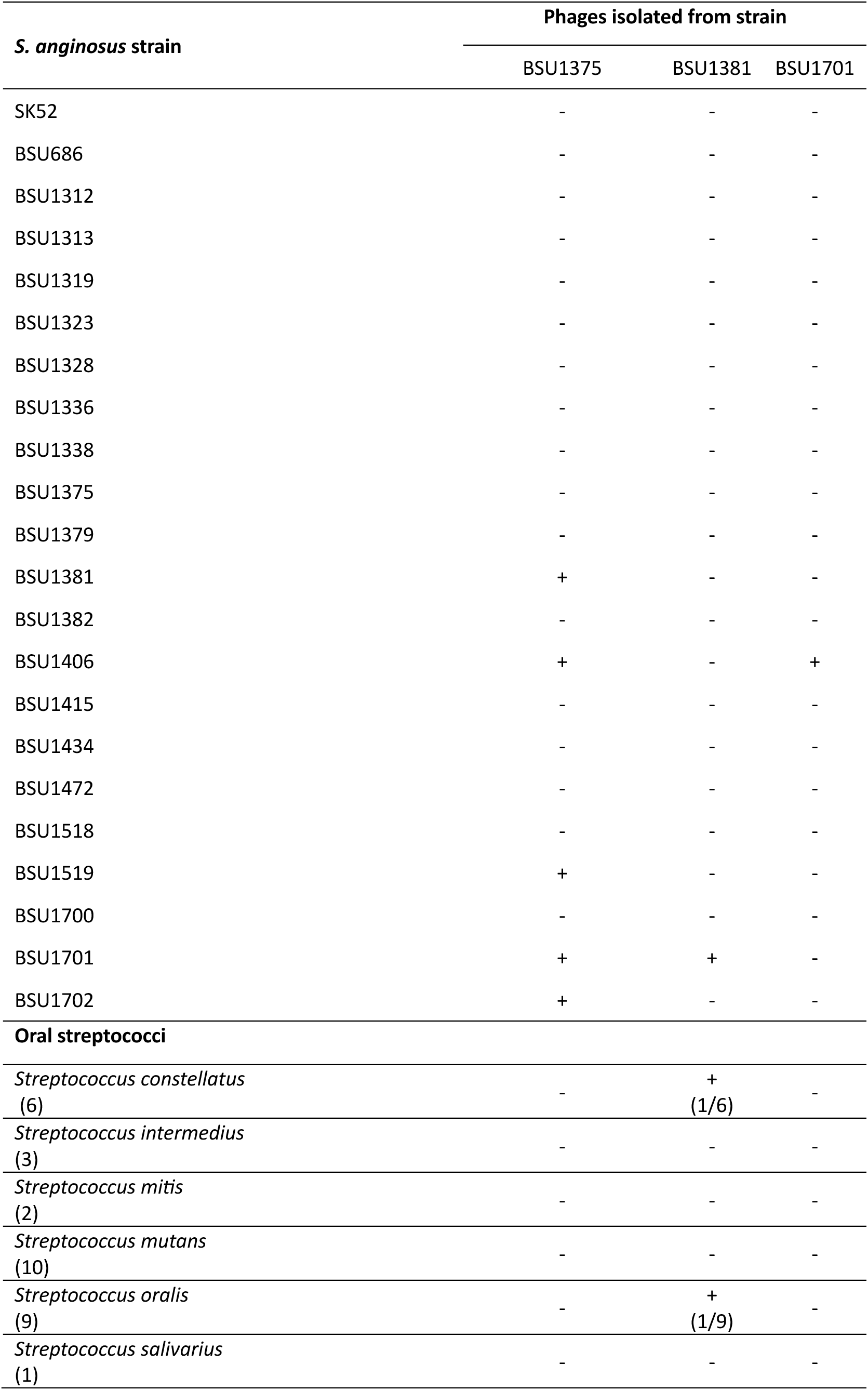
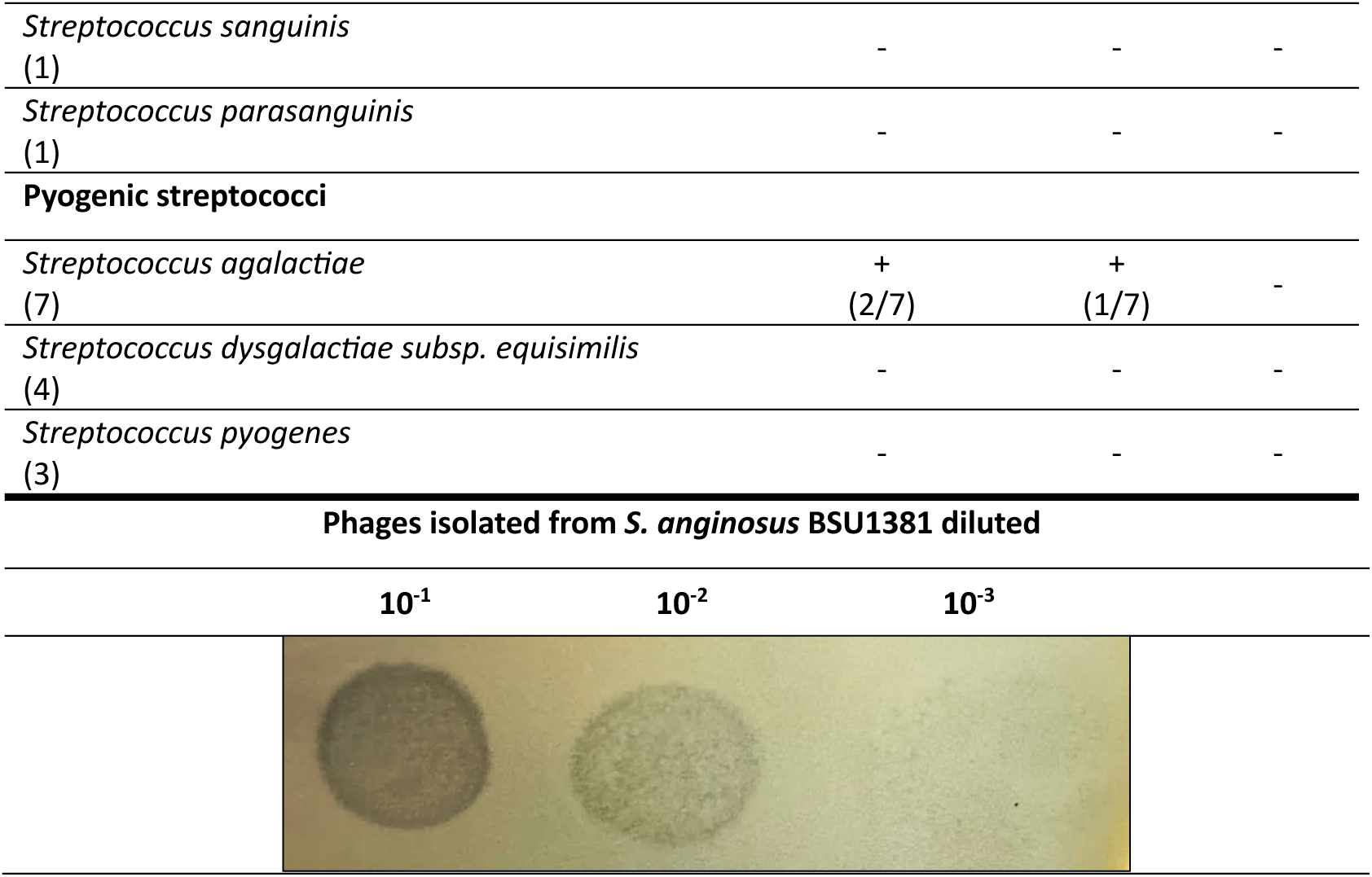
Host range of *S. anginosus* phages. Sensitivity of *S. anginosus*, oral and pyogenic streptococci to isolated phages was assessed in cross-spotting assays. Observed lysis (+) or resistance (-) is indicated. For oral and pyogenic streptococci, the number of isolates tested, and the number of susceptible isolates is noted. Concentration-dependent clearance of the *S. anginosus* BSU1701 bacterial lawn is shown in the lower part. Dilutions of phages isolated from *S. anginosus* BSU1381 were tested in cross-spotting assays.

## Discussion

*S. anginosus* has long been considered a commensal organism of human microbiota, but growing evidence of its involvement in invasive infections has redefined it as an important opportunistic pathogen (Clarridge et al., 2001; Jiang et al., 2020; Laupland et al., 2018; Terzi et al., 2016). Beyond its clinical relevance, recent genomic studies have shown that it contains a remarkable abundance of mobile genetic elements, including integrated prophages (Brassil et al., 2020; Lin et al., 2025; Wang et al., 2025). Prophages contribute significantly to genomic diversity and may impact evolution, virulence, antibiotic resistance, and horizontal gene transfer of their host (Brüssow et al., 2004; Fortier & Sekulovic, 2013; Koskella & Brockhurst, 2014; Schroven et al., 2021). *S. anginosus* prophages have previously been described in genetic studies of strains originating from the urogenital tract (Brassil et al., 2020; Rezaei Javan et al., 2019), but detailed functional characterization is lacking. In the current study we investigated the prevalence of prophages in a collection of 140 *S. anginosus* strains, studied spontaneous and fluoroquinolone-induced prophage excision, examined phage morphology and explored their host range.

To assess the prophage content of *S. anginosus* strains in our collection, WGS data from a subset of strains were analyzed using PHASTEST, whereas the remaining strains were screened by PCR using primers targeting prophage-specific sequences (Brassil et al., 2020). This analysis revealed 31.4% of the 140 strains as lysogenic (Figure 1 and Figure S1), mostly harboring a single prophage. The prophage prevalence of 31.4% in *S. anginosus* appeared lower compared to other streptococci, in which lysogeny was found with varying frequencies among different species (Rezaei Javan et al., 2019). In *S. agalactiae* strains for example, approximately 76% and 84% carry at least one prophage (Shabayek et al., 2025; Wiafe-Kwakye et al., 2024). The presence of prophages in bacterial genomes may reflect a balance between the acquisition of potentially beneficial genes associated with maintaining the lysogenic state and the risk of bacterial lysis following induction of prophages (Bossi et al., 2003; Wendling et al., 2021). Prophage uptake may also be controlled through adaptive immunity provided by CRISPR-Cas systems (Barrangou et al., 2007). Exploring this aspect in our *S. anginosus* strain collection, we found a negative correlation between prophage carriage and the concurrent presence of a type II-A CRISPR-Cas system (Table 2). The observed inverse correlation between prophage carriage and type II-A CRISPR-Cas systems in *S. anginosus* isolates was consistent with findings in other bacterial species. In *S. agalactiae,* a significant difference in prophage content was observed between strains carrying and lacking type II- A CRISPR-Cas (Yamada et al., 2019). Although this correlation is similar in *Escherichia coli* and *Pseudomonas aeruginosa*, it does not appear to be universally applicable, as *Lactobacillus* strains with type II-A CRISPR-Cas systems did not significantly restrict the integration of prophages (Edgar & Qimron, 2010; Pei et al., 2021; Wheatley & MacLean, 2021). However, a major limitation is the PCR-based analysis, which detects only known prophages of *S. anginosus*, likely underestimating the overall prophage diversity. We therefore cannot rule out the presence of novel prophages not yet described in this species. In-depth bioinformatic analysis of our *S. anginosus* strain collection using WGS would be required to explore the full range of prophages, including incomplete or satellite prophages, which may still affect the biology and fitness of bacterial hosts. (Rezaei Javan et al., 2019).

BLAST database searches found all predicted prophages from the *S. anginosus* isolates BSU1375, BSU1381 and BSU1701 to demonstrate close sequence similarity to phages from *S. agalactiae* and *S. anginosus* (Table 1) belonging to the family of siphoviruses. These are the most common phages associated with streptococci (McShan et al., 2019). Siphoviruses are double-stranded DNA viruses with a tail (*Caudoviricetes*), that are characterized by icosahedral capsids, long, flexible, striated and non- contractile tails, as well as a base plate required for recognition of the bacterial host and infection initiation (Kovacec et al., 2024; Malke, 1970; McShan et al., 2019; Sciara et al., 2010). In these experiments, all *S. anginosus* phages exhibited a capsid of about 56 nm (Figure 4). 209 nm long tails and a base plate could be detected in some of the pictures, corresponding well to the typical morphology of siphoviruses (Ackermann, 2007). Overall, the identified structures closely resemble previously described phages of *Streptococcus iniae*, *S. pyogenes*, and *Streptococcus thermophilus* (*S. thermophilus*) and are compatible with the identified nucleotide sequences (Accolas & Spillmann, 1979; Brassil et al., 2020; Harhala et al., 2018; Wright et al., 2013).

Further experiments were conducted to investigate the release and host range of the identified *S. anginosus* phages. Reproducible spontaneous prophage induction was observed for both prophages of BSU1406 as well as for prophage 1375_P1 by PCR. However, prophage 1375_P2 could not be detected in culture supernatants, suggesting that spontaneous prophage induction does not always occur or may not reach detectable levels, as previously described (Czyz et al., 2001; Little & Michalowski, 2010). To increase phage yield, bacterial cultures were exposed to subinhibitory concentrations of fluoroquinolones. Both antibiotics induced phage release well above basal spontaneous levels (Figure 2) and caused concentration-dependent growth differences between prophage carriers and non-prophage carriers (Figure 3 and Figure S2). In conclusion, lysogenic induction by extrinsic factors was successfully established and demonstrated for the first time in *S. anginosus*.

The transition from a dormant, integrated state to the lytic cycle can occur spontaneously or in response to external triggers such as DNA damage (Little & Michalowski, 2010; Lwoff, 1953; Roberts & Roberts, 1975), as has also been observed in other streptococcal species (Baldwin & McKay, 1987; Carrolo et al., 2010). Prophages of the species *S. anginosus* have previously been shown to undergo spontaneous induction (Brassil et al., 2020), although the underlying mechanism that triggers the spontaneous activity of these lysogenic phages may vary (Nanda et al., 2015). Spontaneous phage release is presumed to occur at a limited frequency, yielding phage titers that are several orders of magnitude lower than those obtained after exposure to an inducing agent (Bossi et al., 2003; Little, 2014; Livny & Friedman, 2004). A minor fraction of bacteria sacrificed through spontaneous, prophage-induced lysis may enhance host virulence and biofilm formation in the surviving bacterial population of human pathogens, as described in the studies below. The spontaneous activation of prophages facilitated DNA release upon host lysis, thereby enhancing biofilm formation within the *S. pneumoniae* population (Carrolo et al., 2010). In *Salmonella* strains, the spontaneous release of phage particles confers a competitive advantage over other bacteria (Bossi et al., 2003).

Besides spontaneous induction, several extrinsic DNA-damaging factors can trigger the excision of prophages from bacterial genomes, including the presence of reactive oxygen species produced during the immune response (Figueroa-Bossi & Bossi, 1999; Łoś et al., 2010) and treatment with antibiotics such as mitomycin C (Pricer & Weissbach, 1964), β-lactams (Maiques et al., 2006), and fluoroquinolones (Bearson & Brunelle, 2015; Cirz et al., 2007; Piddock & Wise, 1987). Widely used as broad-spectrum antibiotics, fluoroquinolones interfere with bacterial DNA replication, cause irreversible double-strand DNA breaks, and lead to bacterial cell death (Drlica et al., 2008; Hooper, 2001). However, the application of fluroquinolones at subinhibitory concentrations can trigger prophage induction, as has been reported for *S. pneumoniae* and *Streptococcus canis* (Ingrey et al., 2003; López et al., 2014), but has not yet been observed in the species *S. anginosus*. Significant differences in growth kinetics in the presence of fluoroquinolones between isolates carrying inducible prophages and the non-lysogenic *S. anginosus* SK52 suggested concentration-dependent bacterial lysis by viral particles, consistent with observations in other species (Ingrey et al., 2003; López et al., 2014; Wright et al., 2013). Upon fluoroquinolone treatment in *E. coli*, activation of the SOS response was demonstrated to trigger prophage-mediated cell lysis and markedly increase Shiga toxin production (Wagner et al., 2001; Zhang et al., 2000). As the Shiga toxin is encoded within a prophage, this further underscores the clinical significance of antibiotic-induced prophage excision (Zhang et al., 2000).

The host range of a given phage is primarily determined by specific molecular interactions between phage’s receptor-binding proteins and their receptors on the bacterial cell surface (De Jonge et al., 2019). Although the host range of phages is often strain-specific or limited to closely related species and is generally considered narrow (Brock, 1964; Szymczak et al., 2019), there is growing evidence that it may be broader than previously assumed (De Jonge et al., 2019; Göller et al., 2021; Pchelin et al., 2024; Ross et al., 2016; Whittard et al., 2021). To date, only the purified endolysin of the *S. gordonii* phage PH15 has been described to be capable of infecting *S. anginosus* (Van Der Ploeg, 2008). In this study, however, we were able to demonstrate that fluoroquinolone-induced phages from *S. anginosus* can infect various clinical isolates of this species as well as pyogenic and oral streptococci. Detailed characterization of the *S. anginosus* phages, with particular emphasis on the structural diversity of tail-associated proteins and baseplate components, would provide valuable insights into their biology and the molecular factors underlying host specificity (Bebeacua et al., 2010; Duplessis & Moineau, 2001). The presence of multiple structural proteins mediating the recognition of host receptors has been associated with an expanded host range in *S. thermophilus* phages compared to phages that possess only a single receptor-binding protein (Desiere et al., 1999; Duplessis et al., 2006; Duplessis & Moineau, 2001).

In human hosts, antibiotics such as fluoroquinolones may promote phage-mediated horizontal gene transfer through induction and excision of prophages. Consequently, the spread of virulence and antimicrobial resistance may be amplified, potentially shaping the structure and dynamics of microbial communities, as has been extensively studied in the context of the gut microbiota (Henrot & Petit, 2022) and in enterohemorrhagic *E. coli* (Wagner et al., 2002; Zhang et al., 2000). In addition, also in *Staphylococcus aureus* an association between prophage induction and an enhanced expression of the phage-encoded staphylokinase *sak* was observed upon ciprofloxacin treatment (Goerke et al., 2006). A deeper understanding of prophage induction in *S. anginosus* and other human pathogens may be essential for the development of therapeutic strategies that minimize the unintended activation of prophage-encoded virulence genes and their potential impact.

The oral and pyogenic streptococci analyzed in this study colonize similar niches in the human body as *S. anginosus,* particularly within the oral cavity and the urogenital tract (Olson et al., 2013). The observed infections across different species suggest that the isolated phages possess a broader host spectrum, potentially facilitated by structural similarities among the bacterial surface receptors involved in phage recognition and attachment. Notably, the observed infectivity against *S. agalactiae* may also reflect the partially genetic relatedness of these phages to known *S. agalactiae* phages, suggesting adaptation to similar host-specific characteristics. Furthermore, cross-species infections or, more generally, the exchange of genetic material mediated by prophages may have a crucial impact on the horizontal dissemination of virulence factors between different bacterial species.

This study successfully demonstrated that *S. anginosus* contains various prophages that can be released by spontaneous and fluoroquinolone-triggered excision. The prevalence of prophages seems to be influenced by the presence of type II-A CRISPR-Cas systems in this species, which supports their role in limiting horizontal gene transfer. Visualized *S. anginosus* phages displayed the characteristic morphology of siphoviruses as well as a remarkably broad host range, infecting multiple species of oral and pyogenic streptococci. Taken together, these findings emphasize the functional importance of prophages in the pathogenicity and horizontal gene transfer dynamics of *S. anginosus*, while also drawing attention to the clinical implications of antibiotic-induced prophage activation, particularly regarding stability of the microbiome and the development of therapeutic strategies.

## Material and Methods

### Bacterial Strains and Growth Conditions

Bacterial strains used in this study (Table S1) were stored at -70 °C. *S. anginosus* and other streptococcal strains were cultivated in THY broth [Todd-Hewitt Broth (Oxoid) supplemented with 0.5% yeast extract (BD)]. Cultivation on solid media was conducted on sheep blood agar plates (Oxoid, Basingstoke, UK). Bacteria were incubated at 37°C and 5% CO_2_ overnight, unless otherwise stated.

### General DNA techniques

Genomic DNA (GeneElute^TM^ Bacterial Genomic DNA, Sigma-Aldrich, St. Louis, Missouri) was isolated using commercial kits, following the manufacturer’s protocol. Concentration of obtained DNA was measured using the Quant-iT dsDNA Broad-Range (BR) Assay Kit (Invitrogen, Darmstadt, Germany). Polymerase chain reactions (PCR) were performed applying standard protocols for the *Taq* polymerase. Briefly, after initial denaturation at 95 °C for 5 min 30 amplification cycles of 1 min at 95 °C, 30 s at 50 °C, 1-4 min at 72 °C, and a final elongation time of 7 min at 72 °C followed. The exact annealing temperature and elongation time varied depending on predicted primer-annealing temperature and PCR product length (1 min per 1,000 bp). NucleoSpin® Gel and PCR Clean-up Kit (Machery-Nagel, Düren, Germany) were applied to clean up PCR products and subsequent nucleotide sequencing was conducted by Microsynth SeqLab Laboratories (Göttingen, Germany). Whole genome sequences of *S. anginosus* isolates were obtained based on Oxford Nanopore Technologies performed by Microsynth AG (Balgach, Switzerland). The primers used in this study are listed in Table S2. The PCR-based identification of CRISPR-Cas type II-A systems in *S. anginosus* strains was performed as described previously (Bauer et al., 2020, 2023) with primers designed for the two distinct CRISPR-Cas type II-A loci encoded in two different genomic locations. To analyze the *S. anginosus* strain collection for potential prophage regions, primers were used that have previously been published (Brassil et al., 2020).

### Spontaneous induction of *S. anginosus* prophages

To identify spontaneously induced prophages in liquid culture, specific primers to detect *S. anginosus* prophages were used as described in Brassil et al. 2020. 1.5 ml of overnight-grown bacterial culture was centrifuged at 13,000 g for 10 min at 4 °C, and the supernatant was subsequently filtered through a 0.2 µm syringe filter and underwent DNase treatment, using 1 U DNase I according to the manufacturer’s instructions. Lysates were treated with DNase I to remove free DNA, then heat-lysed to release encapsidated phage DNA, which was amplified by phage-specific PCR. Amplification of the 16S rRNA gene was performed to exclude contamination with bacterial DNA.

### Induction assay with fluoroquinolones in subinhibitory concentrations

First, the MIC of both fluoroquinolones CPX and LVX was determined by adjusting *S. anginosus* isolates to McFarland 0.5 in 3 ml 0.9 NaCl and cultivation on Mueller Hinton agar with 5 % horse blood and NAD. After drying for 15 min, a CPX or LVX Liofilchem® MTS^TM^ test strip was centrally placed on the bacteria containing plate. Strain specific MIC was evaluated after 24 h-48 h incubation time.

Following induction with fluoroquinolones by adding CPX or LVX at subinhibitory concentrations (0.625 µg/ml and 0.3125 µg/ml) to bacterial cultures with an OD_600nm_ of 0.1 in THY, lysogenic growth patterns were analyzed. The control culture was grown in unmodified THY broth. During incubation, the optical density was measured hourly over 7 h and after 24 h.

### Induction, isolation and concentration of *S. anginosus* phages

Bacterial suspensions with an OD_600nm_ of 0.1 in THY were treated with fluroquinolones CPX or LVX in subinhibitory concentrations (0.625 µg/ml and 0.3125 µg/ml) and 10 mM CaCl_2_. After 7 h incubation and centrifugation at 4,707 g; 4 °C for 1 h, the supernatant was filtered through a 0.2 µm syringe filter or Nalgene^TM^ Rapid-Flow^TM^ Filter Unit 0.2 µm with CN membrane. Phages isolated after induction with CPX underwent polyethylene glycol (PEG) precipitation with subsequent chloroform extraction. After incubating the phage lysate with 1 µg/ml DNase I and 1 µg/ml RNase A for 30 min at room temperature, 4 M NaCl was added to a final concentration of 1 M with a subsequent 1 h incubation on ice. Following centrifugation at 4,707 g; 4 °C for 1 h, the supernatant was mixed with 40% PEG_6000_ to a final concentration of 10% and stored on ice overnight. After centrifugation at 4,707 g; 4 °C for 1 h, the dried pellet was resuspended in 400 µl SM buffer (100 nM NaCl; 8 mM MgSO_4_; 50 mM Tris-HCl 1 M pH 7.5) and centrifuged at 17,000 g; 4 °C for 1 h. Dried phage pellet was resuspended in 400 µl SM buffer and subsequently mixed with an equal volume of chloroform. After vortexing for 30 s and centrifuging at 3,000 g; 4 °C for 15 min, the aqueous phase with the concentrated phages, was separated and stored at 4 °C. Phage lysates obtained after treatment with LVX were concentrated by centrifugation at 19,800 g; 4 °C for 1 h and the resulting phage pellets were resuspended in 400 µl SM buffer and stored at 4 °C.

### Transmission electron microscopy

For TEM of concentrated phage samples, suspensions were applied to formvar and carbon coated TEM grids and negatively stained with 2% uranyl acetate as described previously (Olari et al., 2023) and visualized with the transmission electron microscope JEM1400 (Jeol, Tokio, Japan) equipped with a CCD camera (Veleta, Olympus, Tokio, Japan). Measurement of phage dimensions was performed using ImageJ Version 1.53 (Schneider et al., 2012).

### Determination of phage host range

Host range of *S. anginosus* phages was determined by adding 100 µl of bacterial culture to be tested to 3 ml of molten THY soft agar (0.4% agarose) and subsequent spreading onto a Brain Heart Infusion (BHI) agar plate (1.5% agarose). After 15 min, 10 µl of isolated phages were spotted onto bacteria- containing THY soft agar, including a control with 0.625 µg/ml CPX or LVX in SM buffer. Plaque formation was assessed after incubation for 48 h-72 h.

### Bioinformatical and statistical analysis

GenBank database served as source of nucleotide and protein sequences (https://www.ncbi.nlm.nih.gov/). With the PHASTEST tool (Arndt et al., 2016; Wishart et al., 2023; Zhou et al., 2011) bacterial genomes were screened for prophage sequences. Only predicted prophage regions with the completeness level “intact” were considered. Homologous sequences were identified using the Basic Local Alignment Search Tool (BLAST) at the NCBI server (https://blast.ncbi.nlm.nih.gov/Blast.cgi). Genetic analyses were performed with QIAGEN CLC Main Workbench 7 (https://digitalinsights.qiagen.com). The significant association between prophage presence and CRISPR-Cas type II-A presence in *S. anginosus*, was assessed using the Chi-square test of independence using Microsoft Excel. Contingency tables were generated for categorical data, and the CHISQ.TEST function was applied to calculate *p*-values, with statistical significance defined as *p*<0.05. Experimental data is represented as mean value ± standard deviation. The nonparametric Mann-Whitney U test was applied with p-values smaller than 0.05 as significant. Statistical analysis and graph preparation were performed using GraphPad Prism V6.

## Acknowledgements

We thank the staff of the Central Faculty for Electron Microscopy at Ulm University for technical support.

## Author Contribution

BS and DH designed the study. DH, SB, JM and SM performed experiments and analyzed the data. DH and JM did electron microscopy analysis with the support of CR. The manuscript was prepared and edited by DH and BS. CR, PS and GC contributed to the development of the methodology and edited and modified the manuscript. All authors contributed to the article and approved the submitted version.

## Data availability

The raw data supporting the conclusions of this article will be made available by the authors, without undue reservation, to any qualified researcher.

