## Supplementary Tables and Figures for "Fluoroquinolone-Triggered Prophage Induction in *Streptococcus anginosus* Reveals Lytic Cycle, CRISPR–Prophage Interplay, and the potential for Cross-Species Horizontal Gene Transfer"

**Supplementary Figures**

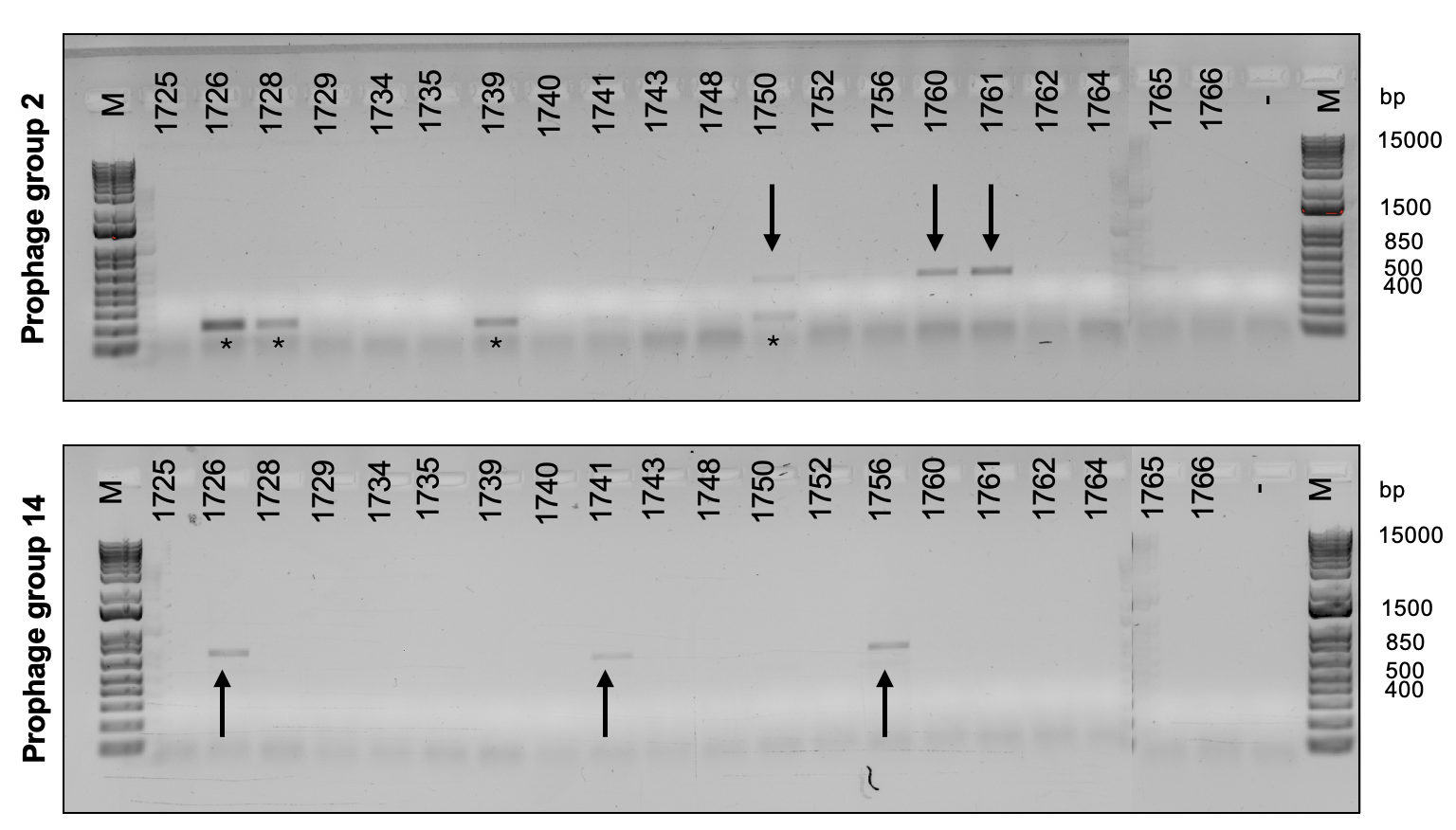

Figure S 1: Prophage screen of S. anginosus isolates. PCR-based analysis with chromosomal DNA as template and primers to detect prophages that have already been published for this species (Brassil et al., 2020). Representative agarose gels are shown. Arrows indicate the viral sequences confirmed by BLAST search, while non-specific fragments are marked with asterisks. The 1 kb Plus DNA Ladder (Invitrogen) served as molecular weight marker (M). Negative control without nucleic acid is labeled with ‘-‘.

Figure S 2: Fluoroquinolone-triggered lysogenic growth patterns of S. anginosus isolates. Growth kinetics of S. anginosus SK52 and clinical isolates BSU1375, BSU1381 and BSU1701 measured after treatment with LVX. Black curves indicate growth without treatment, while 0.625 µg/ml and 0.3125 µg/ml LVX is colored in dark and light red, respectively. Each data point represents an average of five independent measurements including standard deviation. Differences were statistically significant (*p<0.05, **p<0.01, ***p<0.001; Mann-Whitney-U test) and are shown above unless otherwise indicated. When indicated on the left side of the data point it refers to 0.3125 µg/ml and on right side it refers to the untreated control.

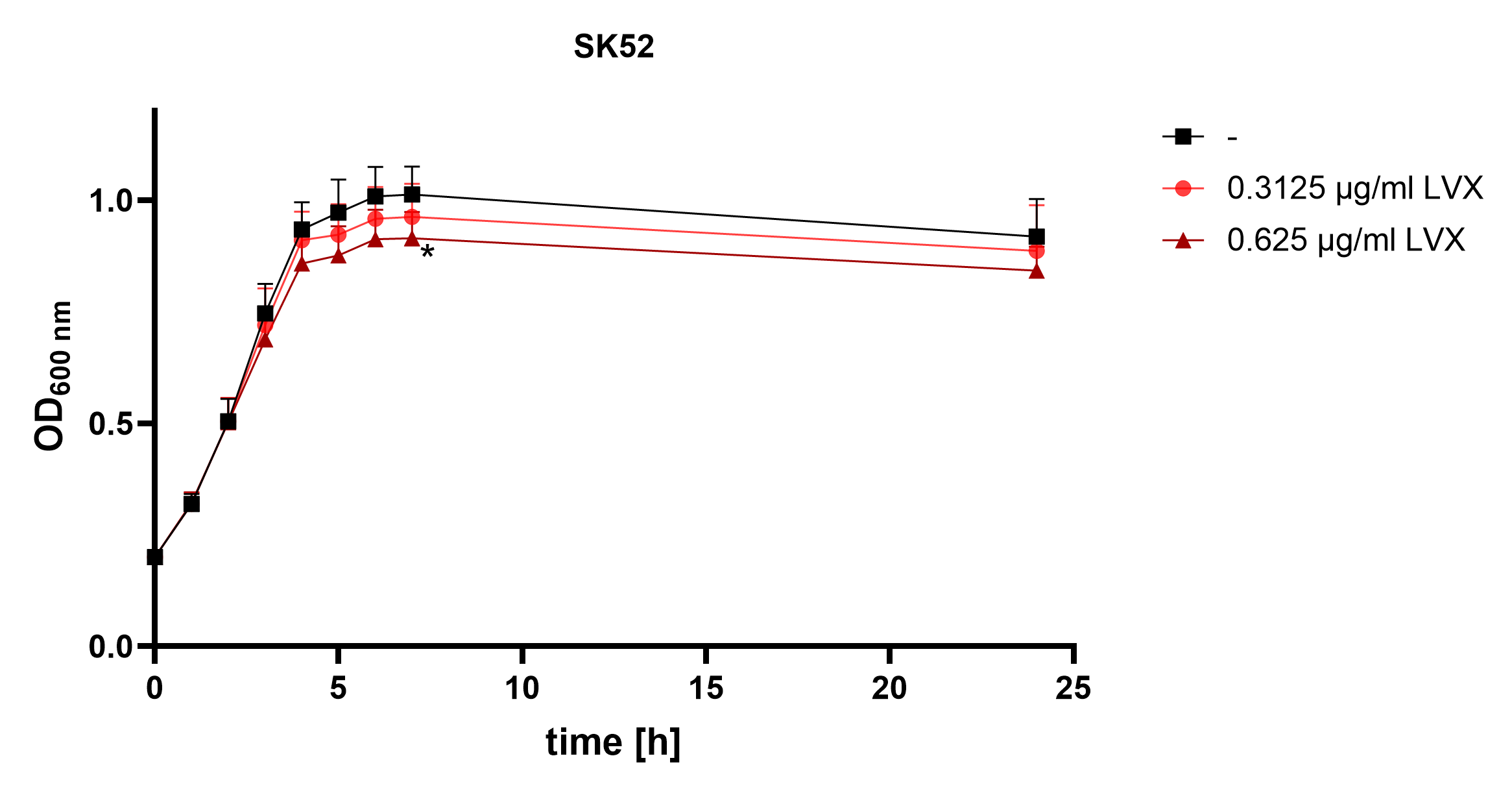

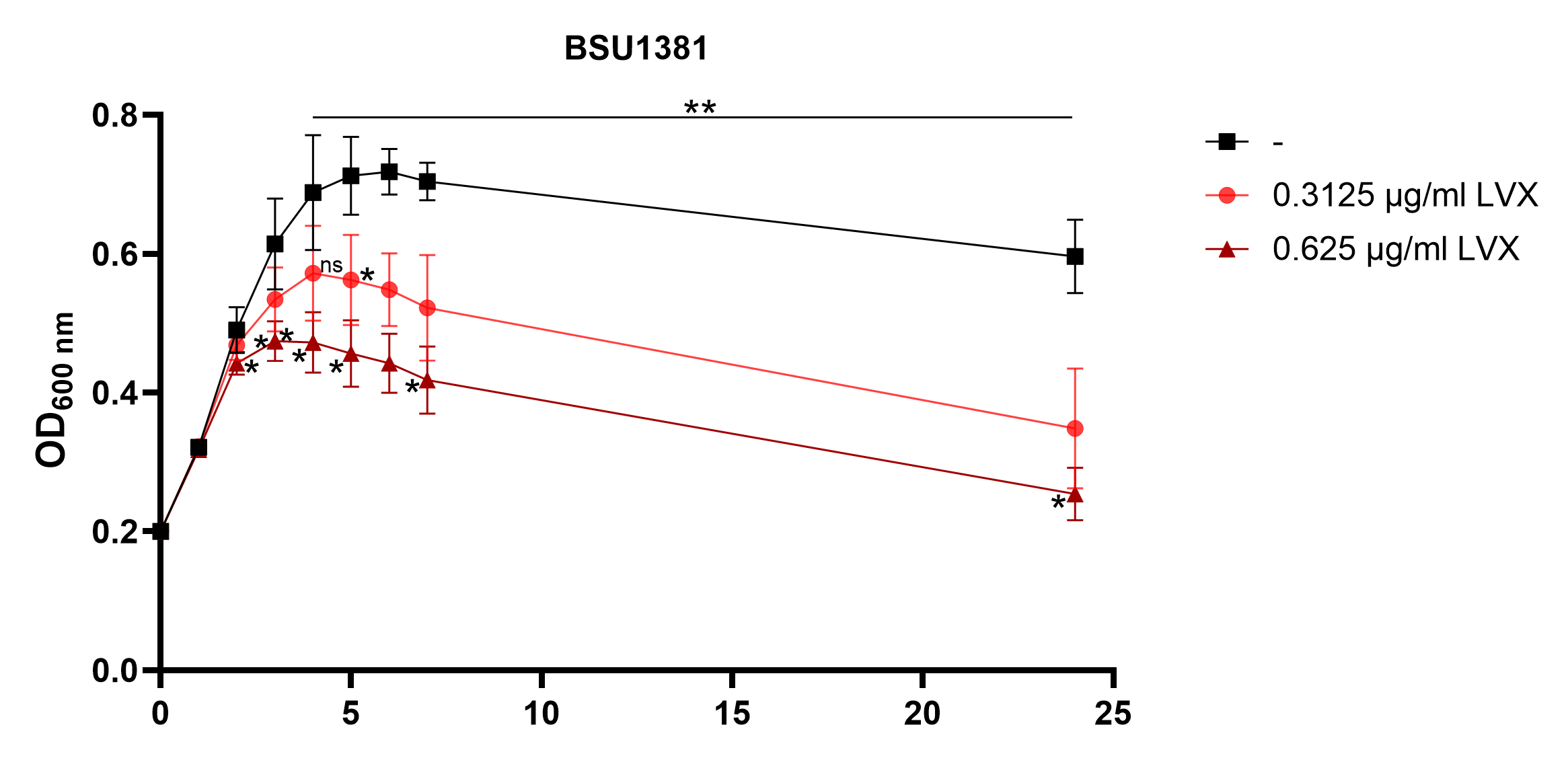

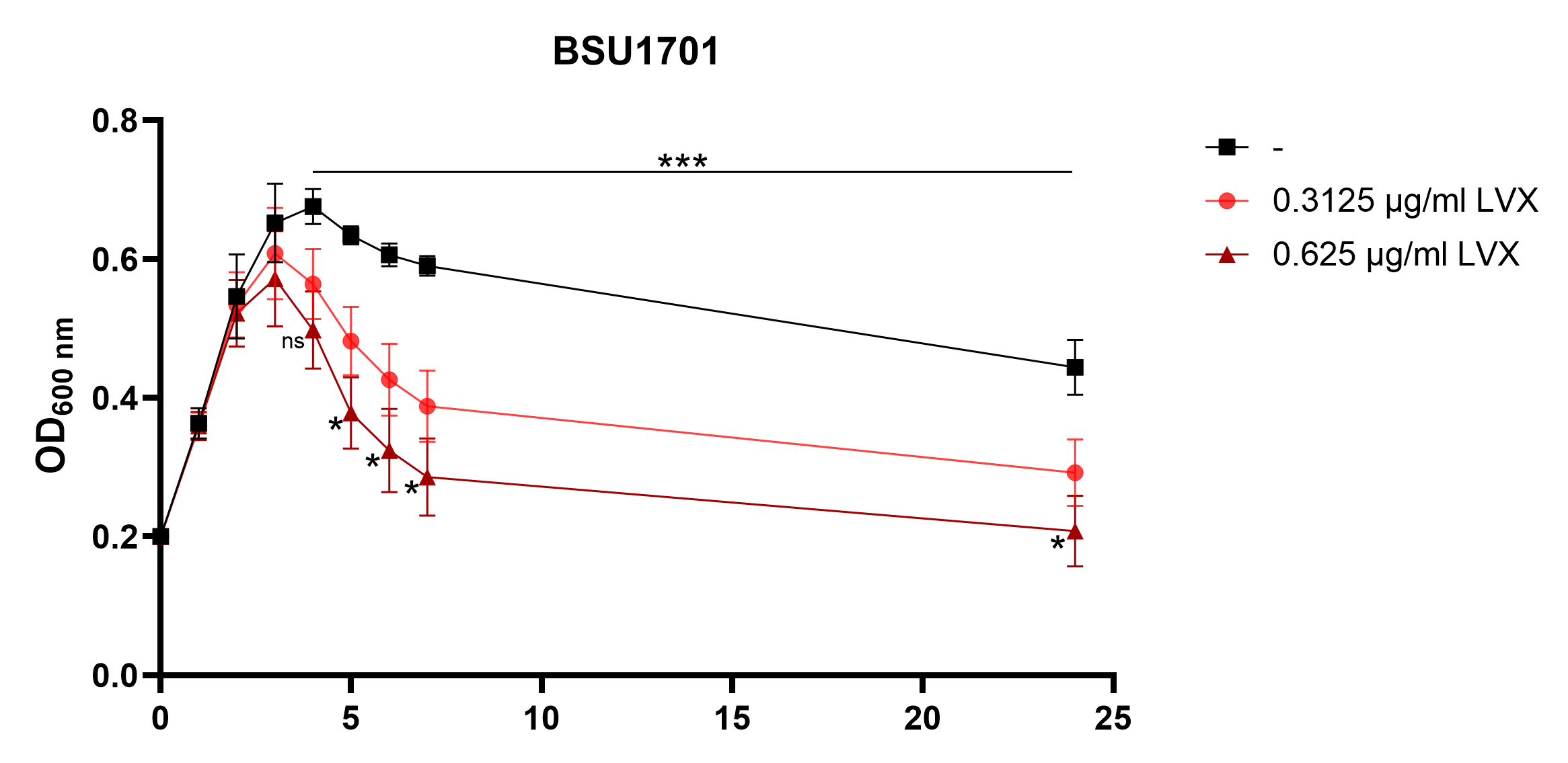

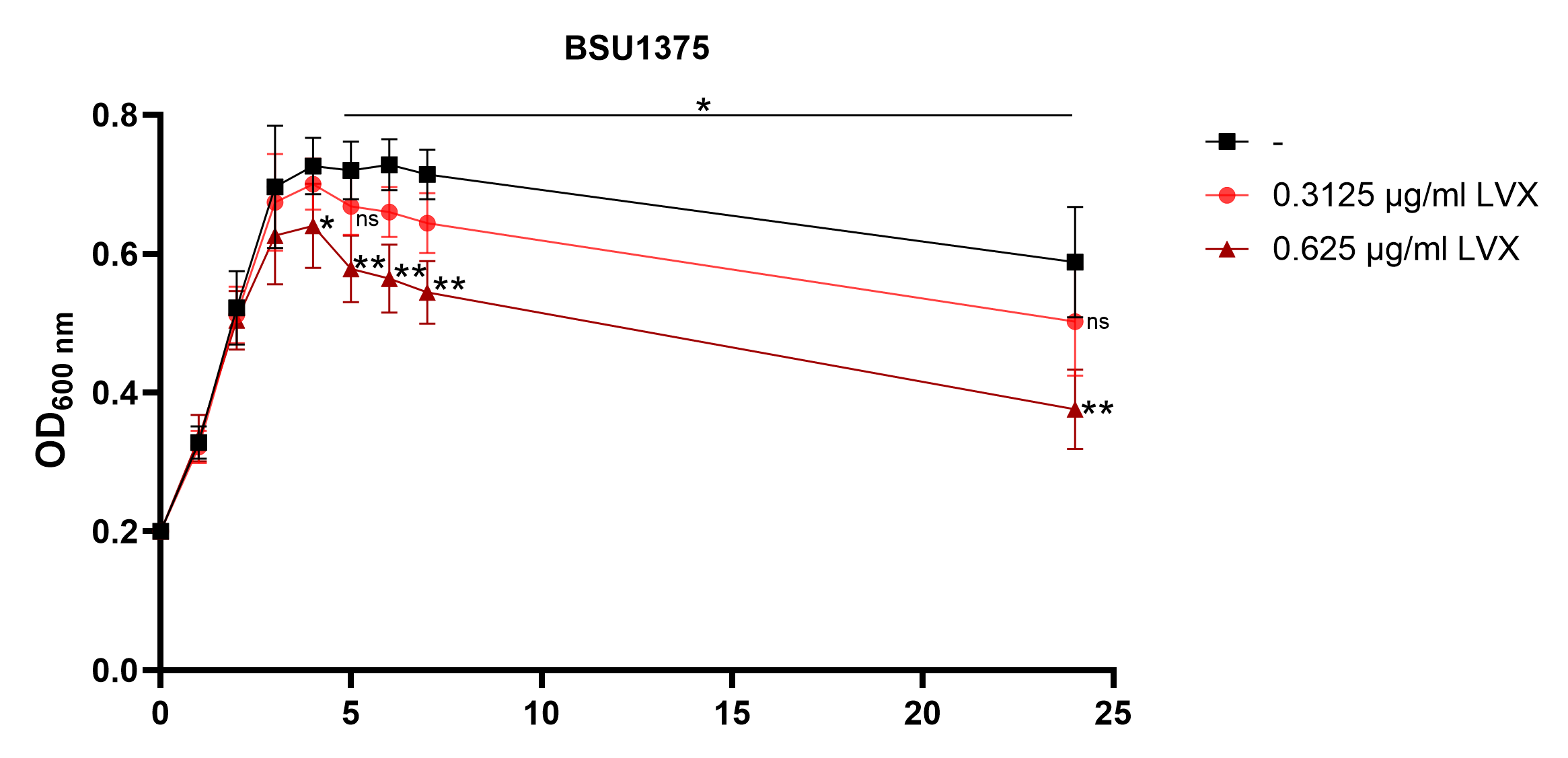

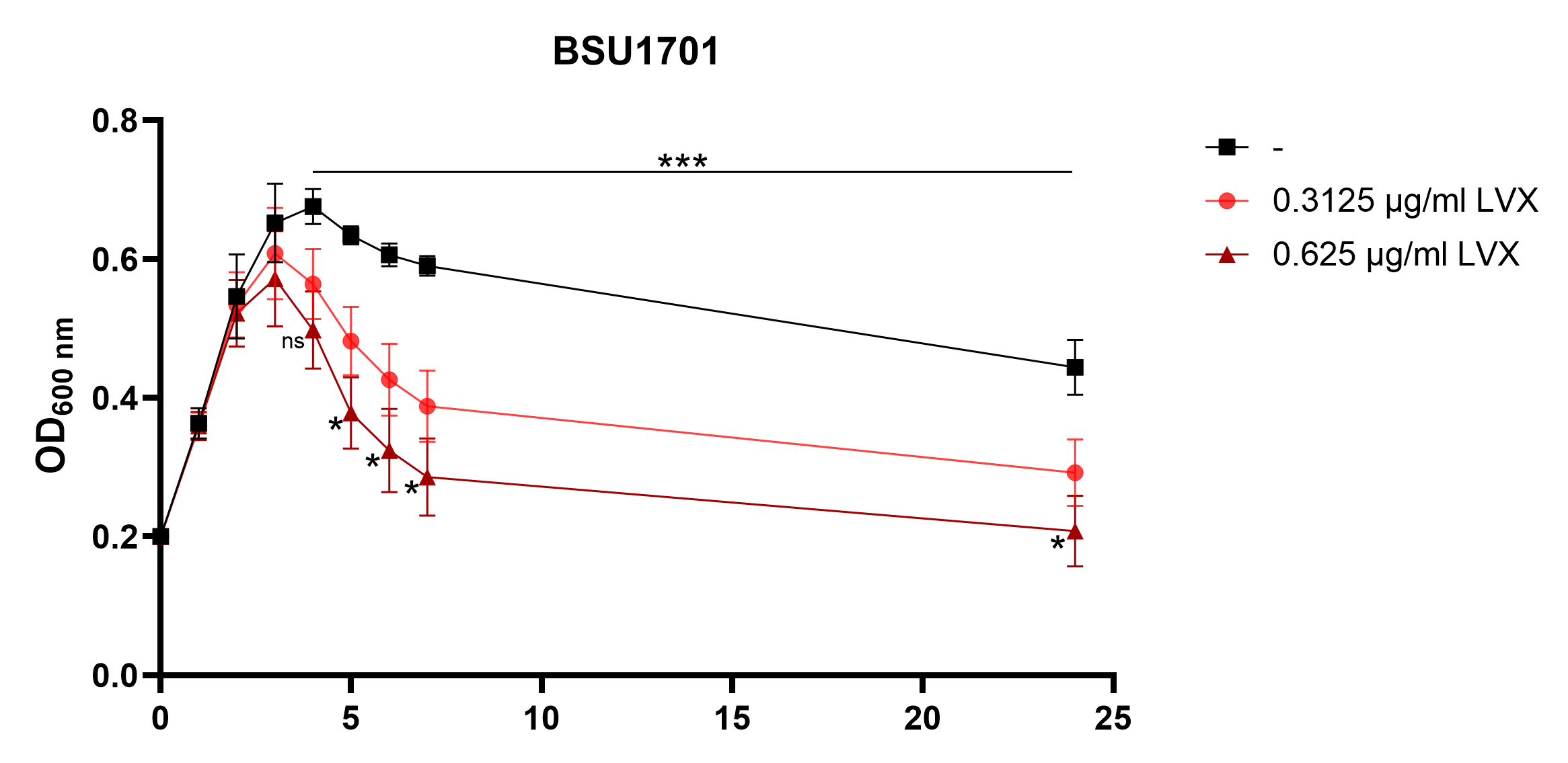

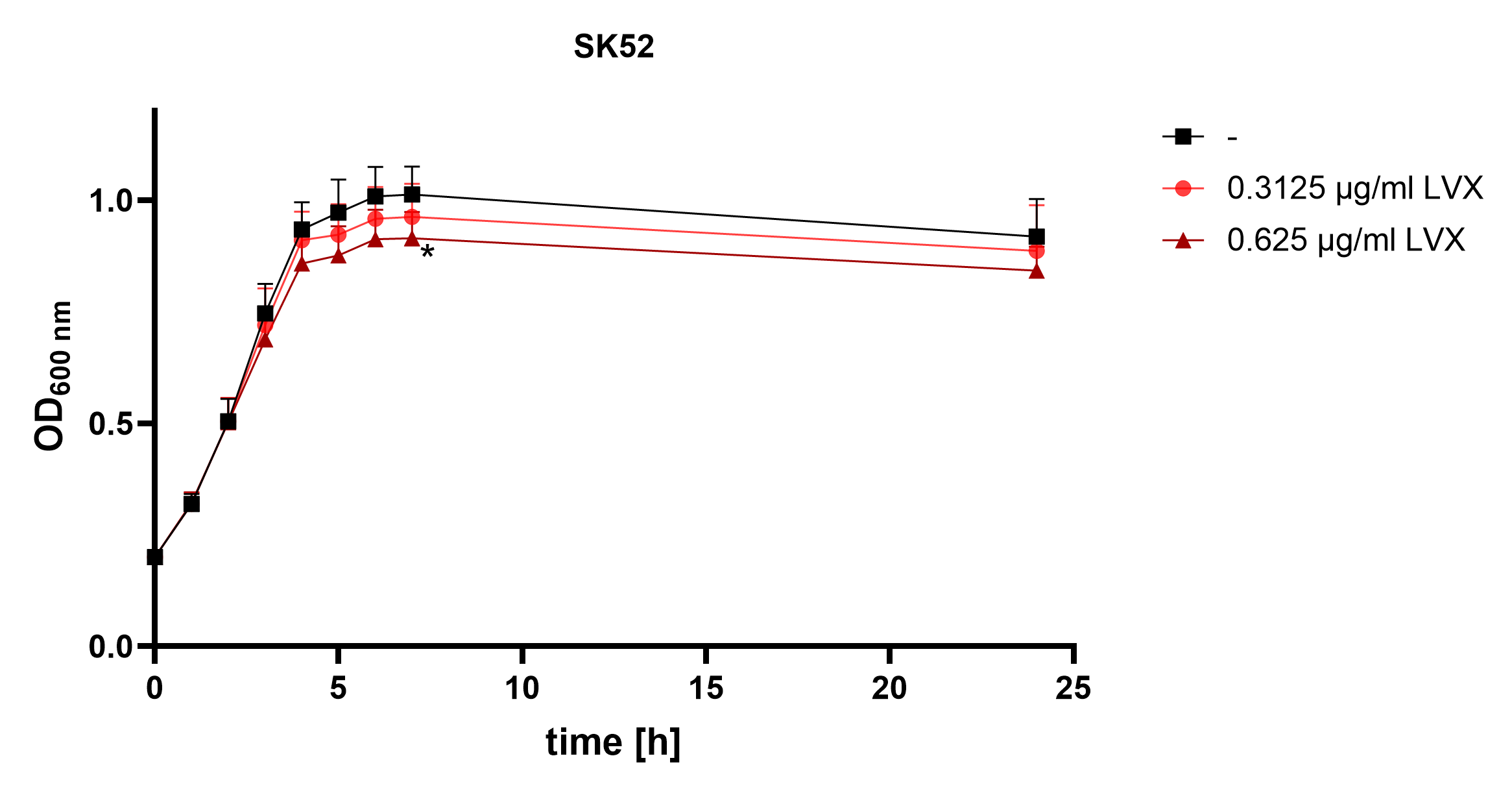

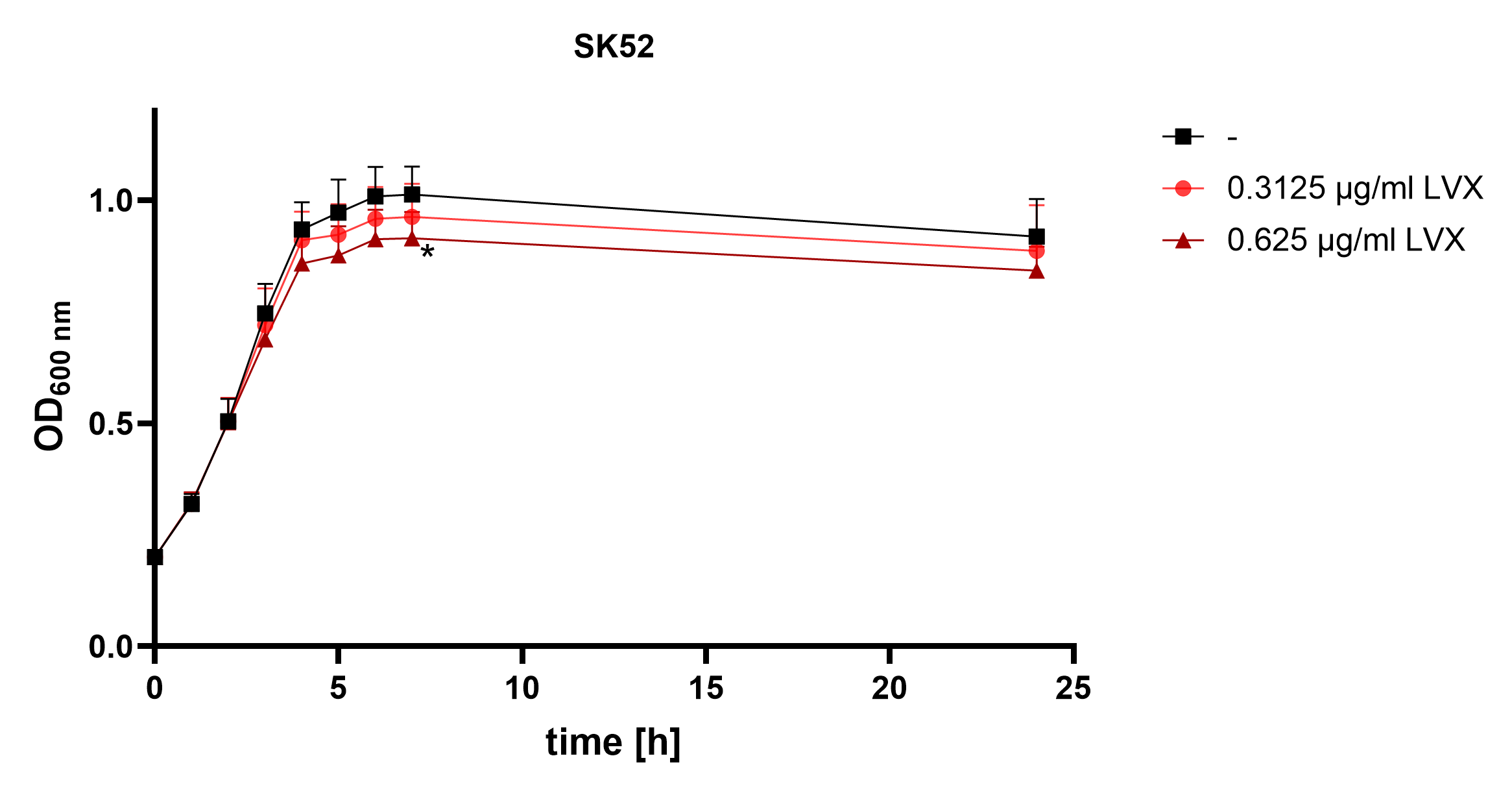

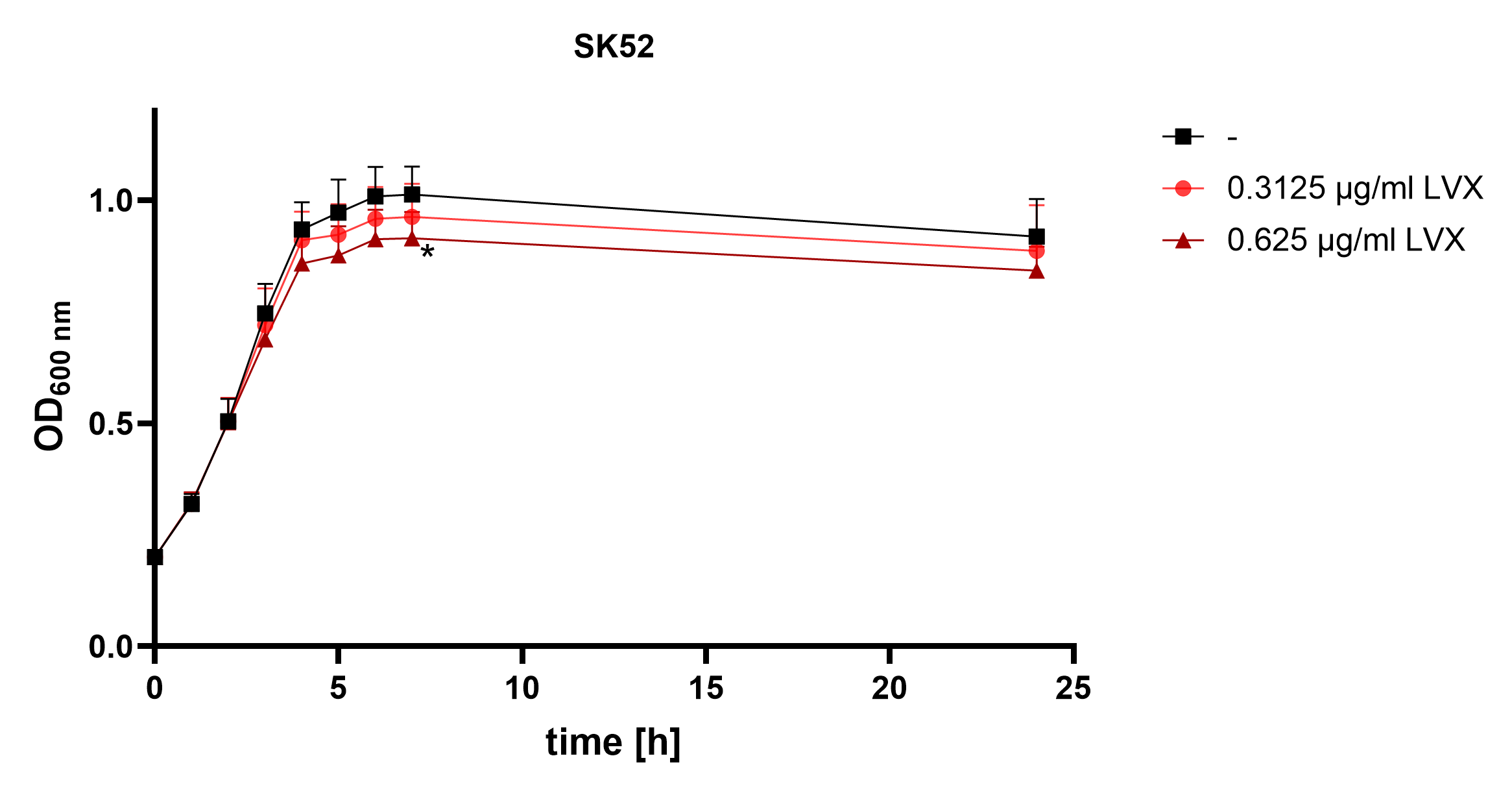

**Supplementary Tables**

Table S 1: Bacterial strains, their definitions and sources.

| Strain | Definition | Source |
| --- | --- | --- |
| Wild type strains |  |  |
| *Streptococcus anginosus* |  |  |
| BSU 458 | *Streptococcus anginosus* type strain SK52, ATCC33397 | ATCC |
| BSU 676, BSU 679, BSU 681, BSU 685, BSU 686, BSU 687, BSU 690, BSU 793, BSU 794, BSU 1210–BSU 1212, BSU 1214–BSU 1217, BSU 1222, BSU 1227, BSU 1289, BSU 1292, BSU 1303, BSU 1304, BSU 1306–BSU 1308, BSU 1310, BSU 1312, BSU 1313, BSU 1317–BSU 1319, BSU 1323, BSU 1324, BSU 1326–BSU 1332, BSU 1334, BSU 1336, BSU 1338, BSU 1339, BSU 1344–BSU 1347, BSU 1349, BSU 1351, BSU 1354–BSU 1356, BSU 1358, BSU 1360–BSU 1363, BSU 1365, BSU 1366, BSU 1369, BSU 1370, BSU 1372, BSU 1373, BSU 1375, BSU 1376, BSU 1379, BSU 1381, BSU 1382, BSU 1384, BSU 1386–BSU 1392, BSU 1395–BSU 1407, BSU 1412–BSU 1415, BSU 1417–BSU 1422, BSU 1434, BSU 1446, BSU 1447, BSU 1473, BSU 1518–BSU 1520, BSU 1793, BSU 1795–BSU 1800, BSU 1802, BSU 1803, BSU 1811–BSU 1813 | clinical isolate | Ulm collection |
| BSU 1448, BSU 1464-BSU 1466, BSU 1472 | clinical isolate^c^ | RWTH Aachen |
| BSU 1698–BSU 1702, BSU 1709, BSU 1714, BSU 1739, BSU 1741, BSU 1743, BSU 1748, BSU 1750, BSU 1752, BSU 1756, BSU 1760, BSU 1761, BSU 1766 | clinical isolate^a^ | Bern University |
| *Streptococcus agalactiae* |  |  |
| BSU 308 | *Streptococcus agalactiae* NEM316, ATCC12403 | ATCC |
| BSU 1023, BSU 1027 | clinical isolate^b^ | Suez Canal University |
| BSU 1301, BSU 1299, BSU 1300, BSU 1511 | clinical isolate | Ulm collection |
| *Streptococcus constellatus* |  |  |
| BSU 459 | *Streptococcus constellatus,* ATCC27823 | ATCC |
| BSU 674, BSU 677, BSU 1290, BSU 1293 | clinical isolate | Ulm collection |
| BSU 1704 | clinical isolate^a^ | Bern University |
| *Streptococcus dysgalactiae subsp. equilimilis* |  |  |
| BSU 238, BSU 263 | clinical isolate^c^ | RWTH Aachen |
| BSU 1218, BSU 1220 | clinical isolate | Ulm collection |
| *Streptococcus intermedius* |  |  |
| BSU 1340, BSU 1352 | clinical isolate | Ulm collection |
| BSU 1703 | clinical isolate^a^ | Bern University |
| *Streptococcus mitis* |  |  |
| BSU 199, BSU 200 | clinical isolate | Ulm collection |
| *Streptococcus mutans* |  |  |
| BSU 268, BSU 673, BSU 875, BSU 1819 | clinical isolate | Ulm collection |
| BSU 1824 | *Streptococcus mutans,* ATCC25175 | ATCC |
| BSU 1825 | *Streptococcus mutans,* NCTC11060^d^ | RWTH Aachen |
| BSU 1826 | *Streptococcus mutans,* KK21^d^ |  |
| BSU 1827 | *Streptococcus mutans,* R254^d^ |  |
| BSU 1828 | *Streptococcus mutans,* R658^d^ |  |
| BSU 1829 | *Streptococcus mutans,* 5DC1^d^ |  |
| *Streptococcus oralis* |  |  |
| BSU 198, BSU 201, BSU 210–BSU 214, BSU 222, BSU 1342 | clinical isolate | Ulm collection |
| *Streptococcus parasanguinis* |  |  |
| BSU 693 | clinical isolate | Ulm collection |
| *Streptococcus pyogenes* |  |  |
| BSU 284 | *Streptococcus pyogenes,* ATCC 12351 | ATCC |
| BSU 998 | *Streptococcus pyogenes,* ATCC 12344 | ATCC |
| BSU 1510 | clinical isolate | Ulm collection |
| *Streptococcus salivarius* |  |  |
| BSU 1000 | *Streptococcus salivarius,* ATCC7073 | ATCC |
| *Streptococcus sanguinis* |  |  |
| BSU 2227 | clinical isolate | Ulm collection |

^a^ Kindly provided by Dr. Parham Sendi, Bern University; ^b^ Kindly provided by Dr. Sarah Shabayek, Suez Canal University; ^c^ Kindly provided by RWTH Aachen University; ^d^ Kindly provided by Univ.-Prof. Dr. Georg Conrads, RWTH Aachen University

Table S 2: Oligonucleotides and their 5’-3’ sequences.

| Primer name | Sequence |
| --- | --- |
| 1406_rev | ACGGGCGGTGTGTACAAG |
| 16s_for | AGAGTTTGATCCTGGCTCAG |
| A_CRISPR_cas9_fwd | TTAAACGGCGTGGTATCAGC |
| A_CRISPR_cas9_rev | ATTCCTGCGCGGTATAAGAC |
| A_CRISPR_control_fwd | GCCTGAAATAATAGTGGTTG |
| A_CRISPR_control_rev | GGTTTTATCATCTTAACTGTC |
| B_CRISPR_cas9_fwd | GGCAGAATATAAGGCGGATG |
| B_CRISPR_cas9_rev | AACCATCATCGATTAGATAG |
| B_CRISPR_control_fwd | AACAGCGCCATCTGGGAAAG |
| B_CRISPR_control_rev | GGTATTAGCGAAGAAGAAGC |
| Phage_gr_0_fwd | TGATCTTGCGTAGGTCAG |
| Phage_gr_0_rev | AGCGCAGACTCAGAGAGG |
| Phage_gr_14_fwd | TCGCTCAATCATCTCATCC |
| Phage_gr_14_rev | GATATGCCGGTCTTGGAG |
| Phage_gr_2_fwd | CTGCAACCTCATCATTGC |
| Phage_gr_2_rev | CCGGCGCTGTCTTATATC |
| Phage_gr_3_fwd | ATTGCGCAAGGACAGC |
| Phage_gr_3_rev | CGAATTGGTGCGACTATG |
| Phage_gr_4_fwd | ATTCGCGCTAAGAAGTGC |
| Phage_gr_4_rev | TGCTCAGAATGTGCTTGG |
